## Supplementary Figures and Tables for "Medium spiny neurons activity reveals the discrete segregation of mouse dorsal striatum"

### Supplementary Information

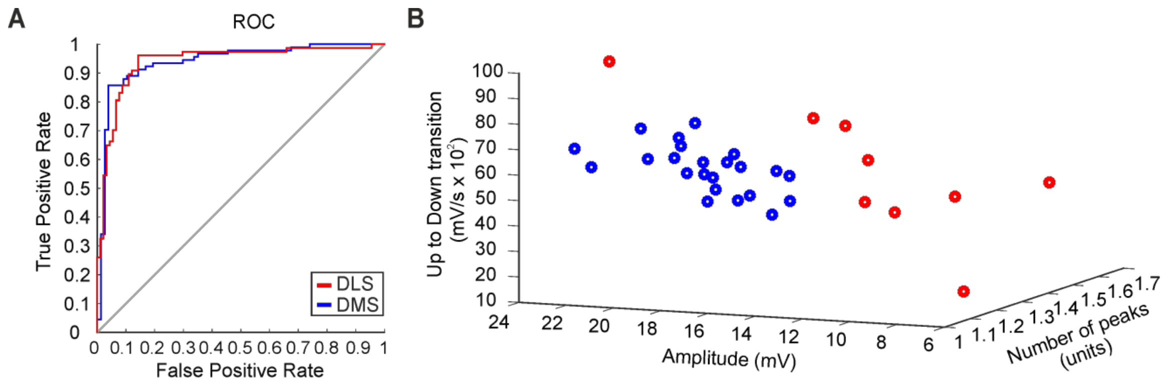


**Figure S1. Classification of DLS- and DMS-MSNs at animal level.**

**A,** One-animal-out classification, ROC curve of the DLS/DMS. **B,** Subspace of classification based on the RFE results. Each dot represents the average of all neurons recorded in single mouse, in DLS (red) or DMS (blue).

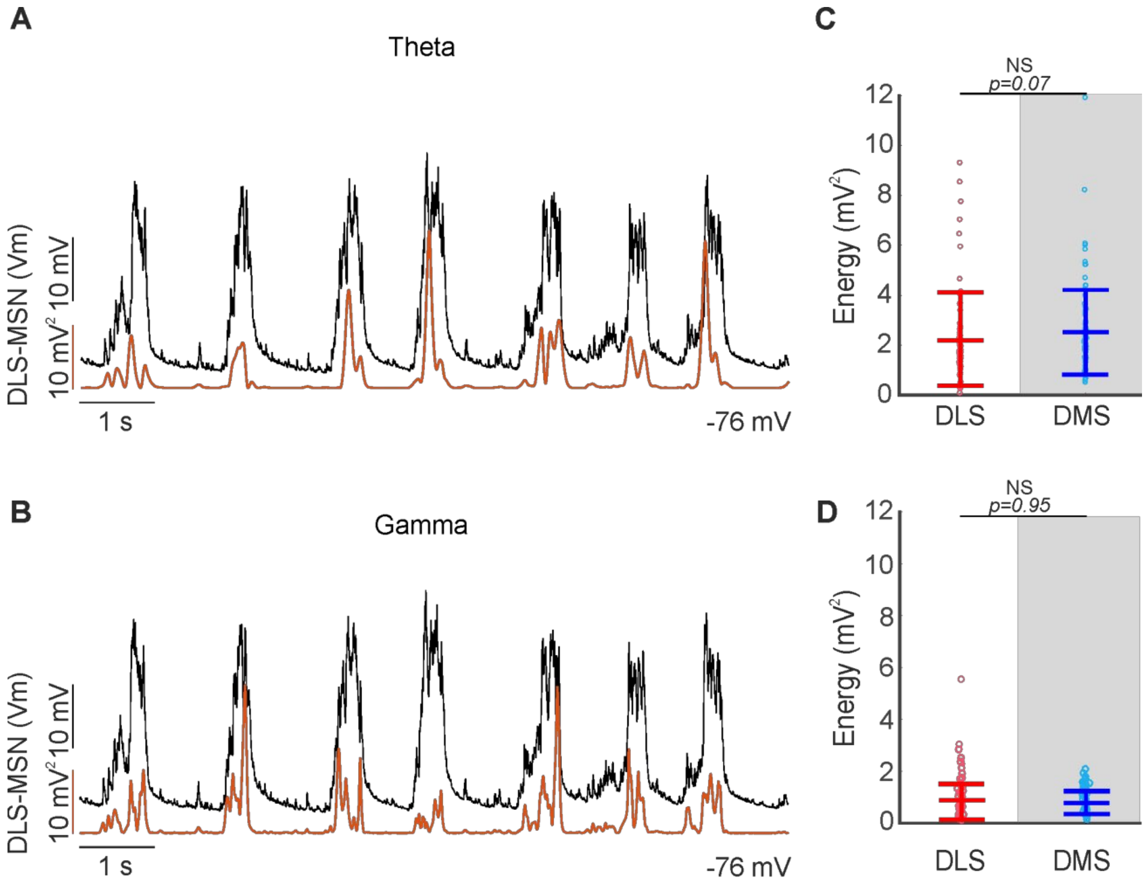


**Figure S2. Theta and Gamma band in membrane voltage of dorsal striatum.**

**A,** Representative example of the energy of the theta band (orange trace) of a DLS-MSN during SWO regime. **B,** Representative example of the energy of the Gamma band (red trace) of a DLS-MSN during SWO regime. **C,** Energy of the Theta band of DLS- and DMS-MSNs. **D,** Energy of the Gamma band of DLS and DMS-MSNs. Same neuron in A and B.

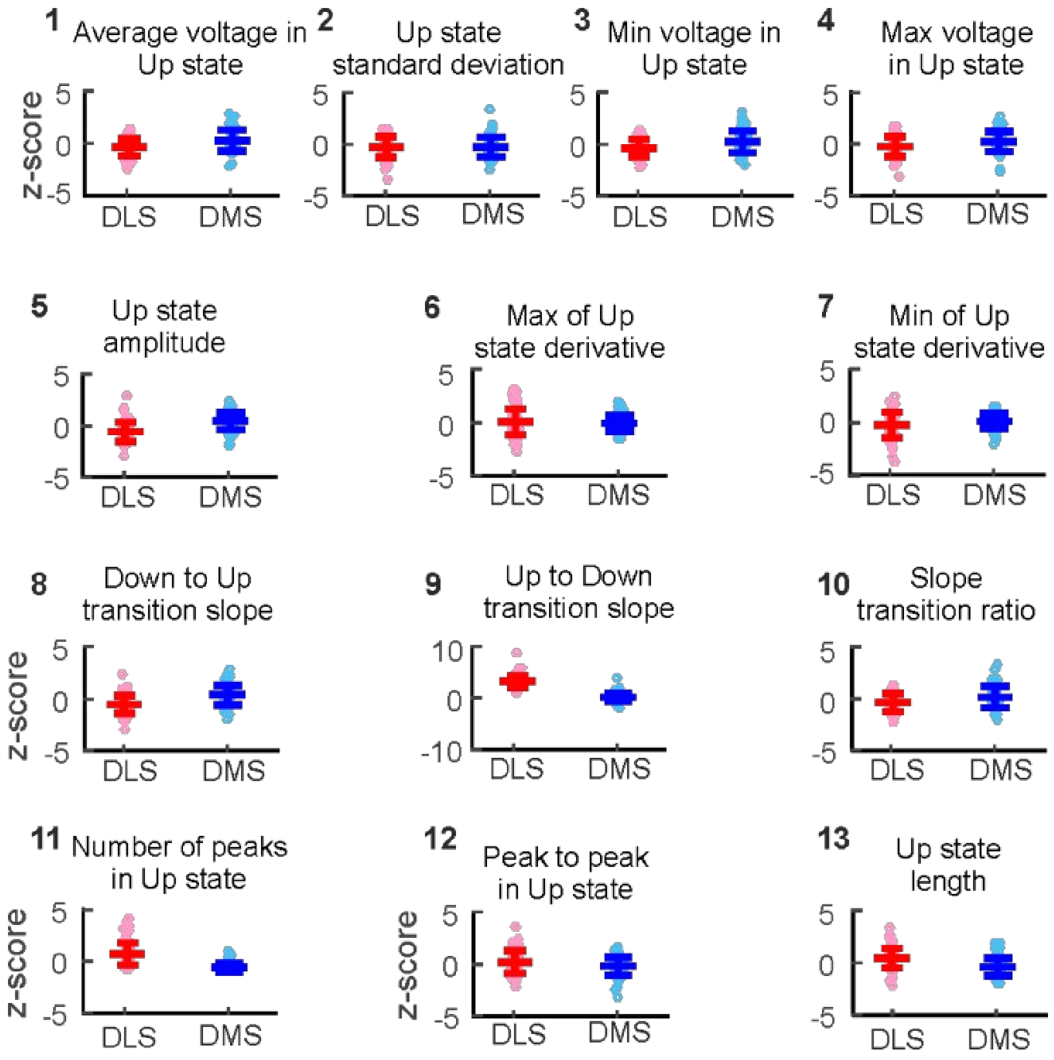


**Figure S3. Z-scoring of the SWO features.**

Z-score transformation of the computed features of the SWO that were used for the classification of DLS- and DMS-MSNs in figure 3A. P values are the same as in figure 3A.

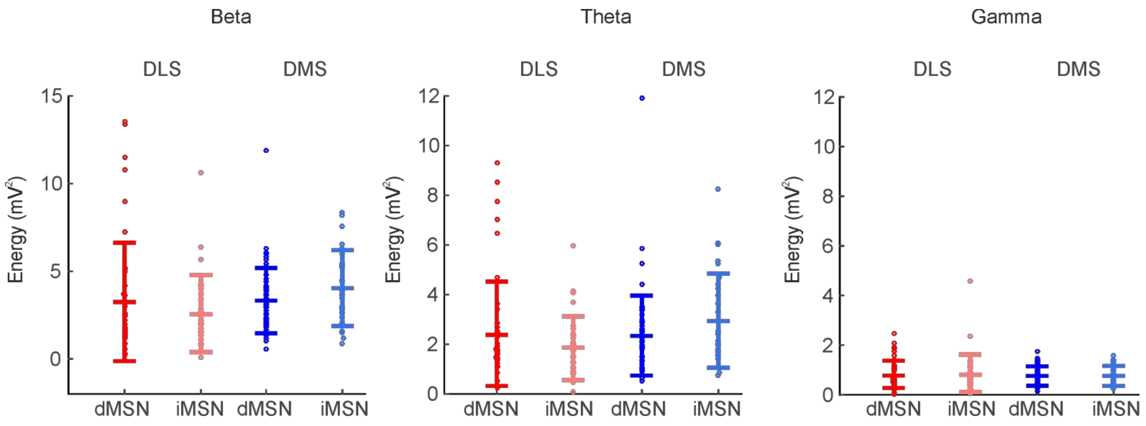


**Figure S4. Beta, Theta and Gamma band in membrane voltage of direct and indirect pathways in the DLS and DMS.** Energy of the Beta, Theta and Gamma band of direct and indirect DLS- and DMS-MSNs.

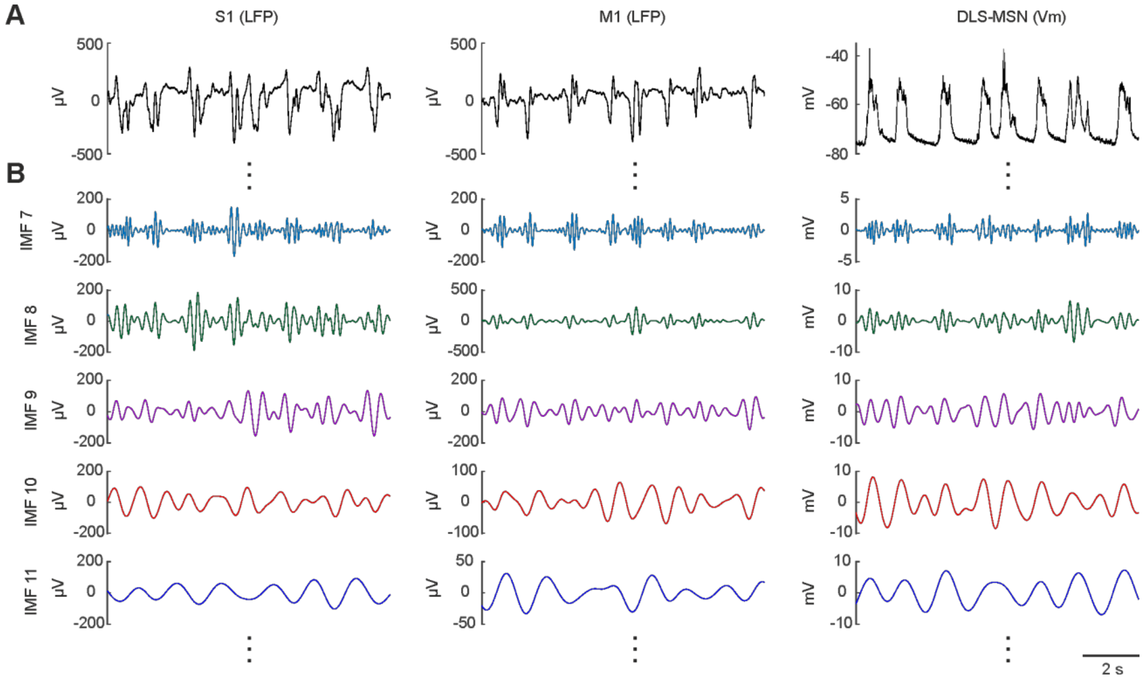


**Figure S5. Example of decomposition of different traces by the NA-MEMD**.

**A,** Example traces of a S1 LFP (left), M1 LFP (center) and a whole-cell recording in DLS (right). **B,** Consecutive IMFs obtained by applying NA-MEMD to the signals in a, starting from the 7$^{th}$ IMF. The IMF classifies the oscillatory activity from the fastest to the slowest.

### Tables

| Feature | Mean value |
|---|---|
| 1- Average voltage in Up state (mV) | -54.44 ± 6.28 |
| 2- Up state standard deviation (mV) | 5.32 ± 1.36 |
| 3- Min voltage in Up state (mV) | -59.95 ± 6.23 |
| 4- Max voltage in Up state (mV) | -43.99 ± 6.20 |
| 5- Up state amplitude (mV) | 13.62 ± 4.10 |
| 6- Max of Up state derivative (dmV/dt) | 0.65 ± 0.14 |
| 7- Min of Up state derivative (dmV/dt) | -0.512 ± 0.12 |
| 8- Down to Up transition slope (mV/s x $10^2$) | 0.829 ± 0.25 |
| 9- Up to Down transition slope (mV/s x $10^2$) | 0.514 ± 0.13 |
| 10- Slope transition ratio (arbitrary units) | 1.61 ± 0.36 |
| 11- Number of peaks in Up state (units) | 1.20 ± 0.15 |
| 12- Peak to peak in Up state (mV) | 20.74 ± 4.65 |
| 13- Up state length (ms) | 431.40 ± 71.50 |

**Table S1. Featurization of the SWO of the DCS-MSNs.**
The number preceding the feature labels are the same as the ones showed in figure 3. All values display means ± standard deviation.

| | |
|---|---|
| **Animals** | 45 |
| **Males** | 19 |
| **Females** | 26 |
| **Weight (gr)** | 33.54±1.37 |
| **Male weight (gr)** | 30.04±1.58 |
| **Female weight (gr)** | 37.58±1.94 |
| **Age (weeks)** | 24.66±1.31 |
| **Male age (weeks)** | 22.52±1.79 |
| **Female age (weeks)** | 27.26±1.84 |
| **Cells/animal** | 4.38±0.49 |
| **Cells/male** | 4±0.92 |
| **Cells/female** | 4.69±0.39 |

**Table S2. Data set description.**
The three first rows show total values, the rest show means ± standard deviation.
